# Climate and vegetation collectively drive soil respiration in montane forest-grassland landscapes of the southern Western Ghats, India

**DOI:** 10.1101/486324

**Authors:** Atul Arvind Joshi, Jayashree Ratnam, Harinandanan Paramjyothi, Mahesh Sankaran

**Affiliations:** National Centre for Biological Sciences (NCBS), Tata Institute of Fundamental Research, GKVK Campus, Bellary Road, Bangalore, Karnataka 560065, India; Manipal Academy of Higher Education, Manipal, Karnataka 576104, India; Research Institute for the Environment and Livelihoods, Charles Darwin University, Darwin NT 0909, Australia; School of Biology, University of Leeds, Leeds LS2 9JT, UK

**Keywords:** Shola-grassland, climate change, soil carbon, land-use, plantations, invasion

## Abstract

Land-use conversion to non-native species plantations not only affects biodiversity but also alters important ecosystem functions including above- and below-ground carbon sequestration, and CO_2_ release rates from soils via soil respiration. Though the role of soil temperature and moisture on soil respiration is well recognized, little is known about how their effects vary across different land-use types. This study looked at the effects of land-cover change on temporal patterns of soil respiration in a montane forest-grassland-plantation matrix, a highly diverse but climatically sensitive ecosystem in the tropical Western Ghats of India. Among native vegetation types, soil respiration rates were higher in grassland compared to forest patches. Invasion of grassland by an exotic tree species - wattle (*Acacia mearnsii*) reduced soil respiration rates to levels similar to that of forests. However, conversion of native grasslands to non-native pine (*Pinus patula*) plantations led to the largest declines in soil respiration rates. In addition, the sensitivity of soil respiration to changes in temperature and moisture differed between different vegetation types. Across all vegetation types, respiration was largely insensitive to changes in soil temperature when moisture levels were low. However, when soil moisture levels were high, respiration increased with temperature in grassland and wattle patches, decreased in the case of pine plantations, and remained largely unchanged in shola forests. Our results suggest that changes in aboveground vegetation type can significantly affect soil C cycling even in the absence of any underlying differences in soil type.

## Introduction

Terrestrial ecosystems, including vegetation and soils, are important components of the global carbon pool, together accounting for about ~2060 Pg of carbon, with about three quarters of this being stored in soils [1,2]. Soils are also primary mediators of global land-atmosphere carbon fluxes, with soil respiration, which results from root respiration and microbial decomposition of organic matter, currently releasing ~75-100 Pg/year back to the atmosphere [3,4]. Like most biochemical processes, rates of soil respiration are largely governed by temperature and moisture availability which play major roles in regulating both plant growth as well as soil microbial activity. Soil respiration rates are typically highest in the tropics where plant growth is luxuriant and conditions for decomposition are optimal, and lowest in cold and dry biomes where microbial activity is limited by low temperatures and moisture availability, respectively [3]. The tropics currently contribute ~67% of the global soil CO_2_ efflux [1,4,5]. Future changes in temperature and precipitation regimes in tropical regions are therefore likely to affect soil respiration rates with potentially significant effects on the global C-cycle and atmospheric CO_2_ levels [1].

Besides climatic variables, changes in land use and land cover also influence global terrestrial-atmosphere CO_2_ fluxes. Since the 1990’s, ~1.14 Pg of C are estimated to have been released back into the atmosphere annually, largely as a result of the conversion of intact forests to secondary forests, pastures and croplands [6–9]. Land-use changes, besides having significant negative impacts on biodiversity, can also alter ecosystem processes by changing the quantity and quality of carbon inputs to soils, changing soil temperature and moisture regimes and by altering hydrological cycles [3, 10–12]. However, the impact of such land-use changes on ecosystem processes, particularly in tropical regions, has received very little attention. More accurate quantification of the effects of land-use changes on carbon fluxes to the atmosphere is needed for better implementing policies such as REDD (Reducing Emissions from Deforestation and Degradation), especially in the tropics which account for the substantial proportion of the emissions due to land-use changes [13].

In this study, we quantified soil CO_2_ effluxes over three years across different land-use types in the climatically sensitive tropical montane forest-grassland mosaics of the Western Ghats biodiversity hotspot in India. These mosaics have undergone extensive land-use changes since the mid-nineteenth century, primarily through the establishment of non-native plantations in grasslands especially of acacia (*Acacia* spp.), eucalyptus (*Eucalyptus* spp.), pine (*Pinus* spp.) and tea (*Camellia sinensis*). Specifically, we looked at how land-cover changes influenced both total and temporal patterns in soil respiration, and how such changes were mediated by alterations to moisture and temperature regimes.

## Methods

### Study area

Our study was conducted in the high elevation forest-grasslands mosaics of the upper Nilgiri landscape (11.27° N 76.55° E, elevation: ~2300 m) in the southern Western Ghats, India (Fig 1). The upper reaches (1200 m to 2650 m) of India’s Western Ghats biodiversity hotspot (8° to 21°N, 73° to 78°E) are characterized by stunted evergreen tree forest patches, locally known as sholas, embedded within grasslands, with abrupt transitions between them (Fig 1). These bi-phasic mosaics are Pleistocene relics that have been in existence for more than 20,000 years [14,15], and support a diverse array of plant and animal species, many of which are endemic [See:16–20].

**Fig 1.**
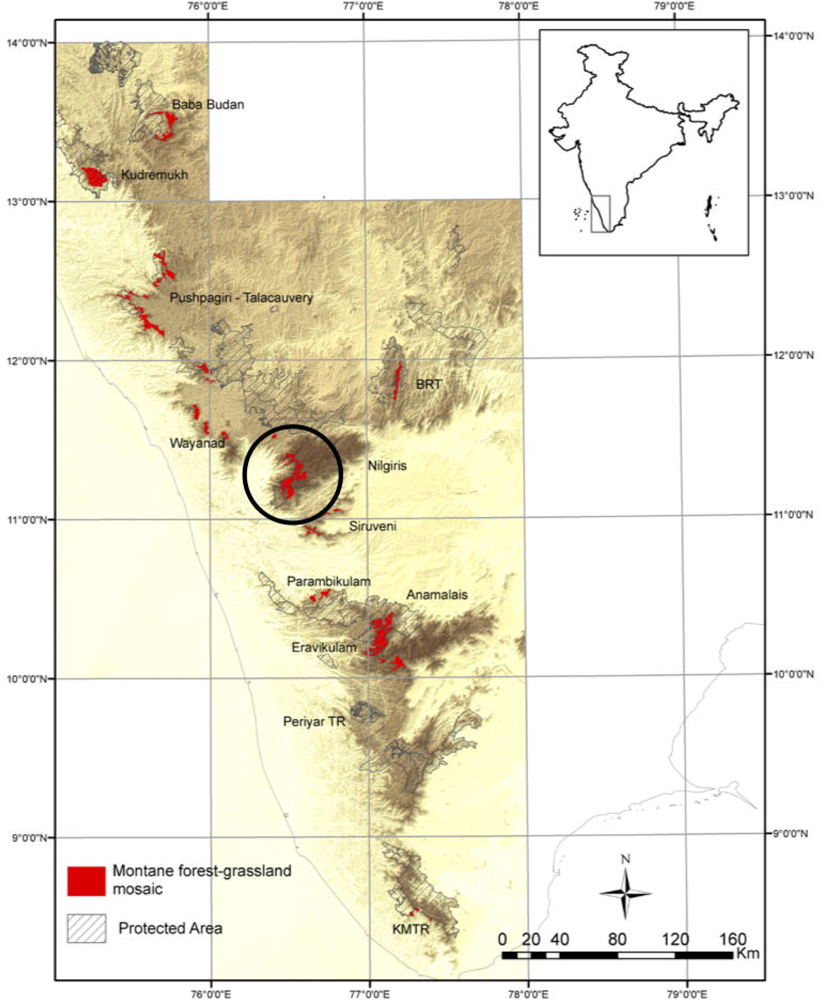
Map showing the distribution of shola-grassland ecosystems across the Western Ghats biodiversity hot-spot, with the study area circled. (Reproduced with modification from Das et al. 2015 [23]).

Large-scale plantations of alien tree species, chiefly of eucalyptus (*Eucalyptusglobulus*), pine (*Pinuspatula*) and wattle (A. *mearnsii*) were established in grasslands of this landscape in the latter half of the nineteenth century [21,22]. Among these, wattle eventually became invasive and now occupies large tracts of former grassland. The landscape is currently characterized by a mix of native and exotic vegetation. The prominent land-cover types include native shola forests, grasslands, invasive wattle patches, and plantations of eucalyptus, pine and commercial tea. We selected four land-use types for this study that were in close proximity to each other - native shola forests, grasslands, invasive wattle patches and pine plantations.

Average annual temperatures in the study area range from a minimum of 5°C in January to a maximum of 24°C in April [24]. Nocturnal frosts are common in grasslands during the winter (November to March) when temperatures routinely drop down below 0°C. Annual precipitation is spatially and temporally variable across the landscape, ranging from 2500 to 5000mm, with most of the rainfall received between June and October from the south-west monsoon. Soils of the area are derived from parent rocks which are gneiss, charnockites and schists [24].

### Study design

In October 2014, we identified three replicate sites in each of four different land-use types: shola forests, grasslands, wattle patchesand pine plantations for the study. Replicate sites within each land-use type were separated by a minimum distance of 100m. At each site, we established five soil respiration collars spaced at least 1m apart from each other. Respiration collars were 10cm long open-ended PVC pipes with an inner diameter of 10cm, inserted 3cm into the soil with 7cm exposed above the soil surface. Collars remained permanently installed in the soil throughout the study, with a few exceptions when collars were lost (due to animal activity or burnt following a fire event) and had to be replaced. In such cases, collars were replaced immediately, and at least a week before the next measurement. In total, we had 60 respiration collars across the different land use categories.

Soil respiration was measured every 2 weeks at each of the 60 collars using an EGM-4 Environmental Gas Monitor (EGM-4, PP Systems, USA). All measurements were made between 10 am and 4 pm. All plant material, primarily litter, within collars was carefully removed before respiration measurements were taken. CO_2_ efflux rates were calculated following Marthews et al. 2014 [25]. Alongside CO_2_ efflux measurements, ambient atmospheric temperature, soil temperature and soil moisture were also quantified. Soil temperature and soil moisture were measured at 0-12 cm depth at three locations around each collar using a handheld long-stem thermometer (HI145-00 and HI145-01, Hanna Instruments, USA) and soil moisture using a FieldScout TDR 200 soil moisture meter (Spectrum Technologies, USA), respectively.

Given the possibility of measurement errors in CO_2_ effluxes while using the EGM in the field, only those CO_2_ efflux measurements which satisfied a criterion of a positive slope with when CO_2_ accumulation was plotted against time, with R^2^ ≥ 0.9, were used in subsequent analyses [26]. Further, we expected soil respiration values to be similar among the five replicate collars within a site. In cases where CO_2_ efflux of an individual respiration collar was less than half or greater than twice the mean of the CO_2_ efflux of the other four adjacent respiration collars at a site, the measurement was removed from the dataset. All data were then averaged by site for the analysis. Annual estimates of soil respiration (g.m^−2^.yr^−1^) for each land-use type were obtained by scaling up the mean hourly CO_2_ efflux rates (g.m^−2^.hr^−1^) estimated for each site for the corresponding year.

We analysed the effects of land-use type, soil temperature and soil moisture on soil respiration using a linear mixed effect model framework with the interaction of land-use type, soil temperature and soil moisture as fixed effects and site as a random effect. Soil respiration values were log transformed before analyses to meet the assumptions of homogeneity and normality of residuals.

The R package nlme with *lme* [27] was used to conduct the mixed effect model analysis. All analyses were conducted using R version 3.3.2 [28].

## Results

### Temporal dynamics of soil respiration across land-uses

Soil respiration rates across land-use types varied both seasonally and annually, and ranged from 0.07 to 2.2 CO_2_ g.m^−2^.hr^−1^ across the study in the different land use types. Soil CO_2_ efflux typically peaked during the south-west monsoon season (June-October), and declined through the winter (November - January) to lowest values in summer (February to May; Fig 2a). Highest respiration rates were recorded in grasslands (average 0.57 g.m^−2^.hr^−1^ across the study) and lowest rates from pine soils (average 0.42 g.m^−2^.hr^−1^, Table 1; Fig 2a). Annual soil CO_2_ effluxes in all land-use types were higher in 2016–17 compared to the previous two years (Fig 3). This increase appears to be associated with a more even distribution of precipitation throughout the year rather than higher total rainfall amounts (Fig 3).

**Fig 2.**
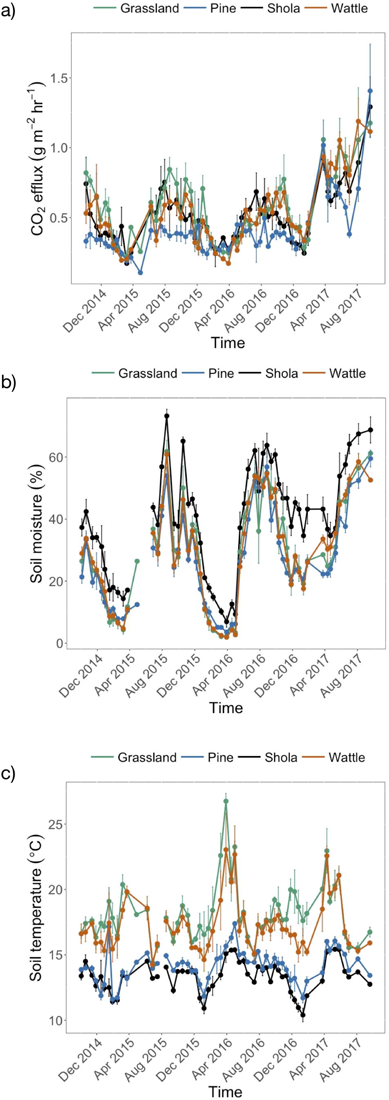
Patterns of change in a) soil respiration, b) soil moisture 0-12cm below the surface and c) soil temperature 0-12 cm below surface over time (n=12). (gaps in 2b and 2c are due to no measurements during the correspondent period because of instrument failure).

**Fig 3.**
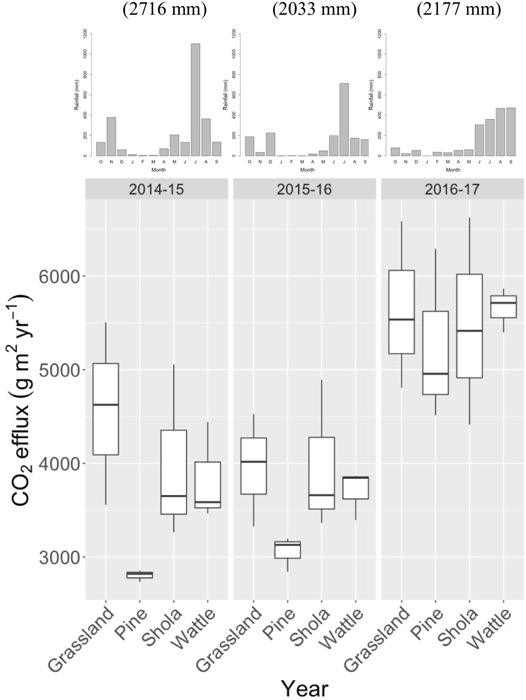
Annual estimates of soil CO_2_ effluxes across different land-use types based on three years of monitoring. Bar graphs on top show monthly precipitation measured at study sites during the corresponding year. Total precipitation during the corresponding year are shown in brackets on top of bar graphs.

**Table 1.** Mean soil temperature, soil moisture and soil respiration across the land-uses. Range presented in brackets

| Land-use category | Soil temperature at 0-12cm depth (°C) | Soil moisture at 0-12cm depth (%) | Soil respiration (CO<sub>2</sub>g.m<sup>-2</sup>.hr<sup>-1</sup>) |
| --- | --- | --- | --- |
| **Grassland** | 18.3 (10.8-31.8) | 29.7 (1.4-70.7) | 0.57 (0.14-2.2) |
| **Pine plantation** | 14.1(10.1-21.6) | 28.2 (1.7-70.3) | 0.42 (0.10-2.2) |
| **Shola forest** | 13.4 (9.2-19.6) | 41.7 (3.4-84.6) | 0.51 (0.07-1.73) |
| **Wattle patches** | 17.4 (11.6-29.0) | 29.7 (1.1-66.8) | 0.52 (0.10-1.9) |

Seasonal soil CO_2_ effluxes mirrored patterns in soil moisture. Over the course of the study, soil moisture levels ranged from 1% to 85%, and not unexpectedly, peaked during the monsoon and were lowest in summer (Fig 2b). Soil moisture levels were higher in shola forests (average 42%) compared to other land-use categories which did not differ from one another (average: 29%; Table 1). Soil temperature was similarly highly variable and ranged from 9.2 to 31.8°C over the course of the study. As expected, soil temperature also showed a distinct seasonal pattern, being highest in summer (April) and lowest during the winter (December) (Fig 2c). Temperatures were consistently higher, but also more variable in grasslands and wattle patches compared to closed canopy shola forests and pine plantations (Table 1, Fig 2c).

### Interactive effects of land-use type, soil temperature and moisture on soil respiration

The linear mixed model analysis revealed a significant interactive effect of land-use type, soil moisture and soil temperature on respiration, collectively explaining 66% of the variation in the data (*F* = 4.48, *df* = 3, *P* = 0.004; Table S1). To visualize this interaction, we modelled soil respiration as a function of temperature for fixed soil moisture levels across the different land use types (Fig 4). Soil moisture values were fixed at the quantiles of measured instantaneous soil temperature measurements over the study (Fig 4). In both grassland and wattle patches, soil respiration was not affected by soil temperatures when moisture was low, but increased with increasing soil temperature at high moisture levels (Fig 4). In pine plantations, respiration was similarly largely unaffected by temperature at low moisture levels, but decreased with increasing temperatures when soil moisture was high (Fig 4). In contrast, soil respiration did not seem to be significantly impacted by changes in either soil temperature or moisture in shola forests (Fig 4).

**Fig 4.**
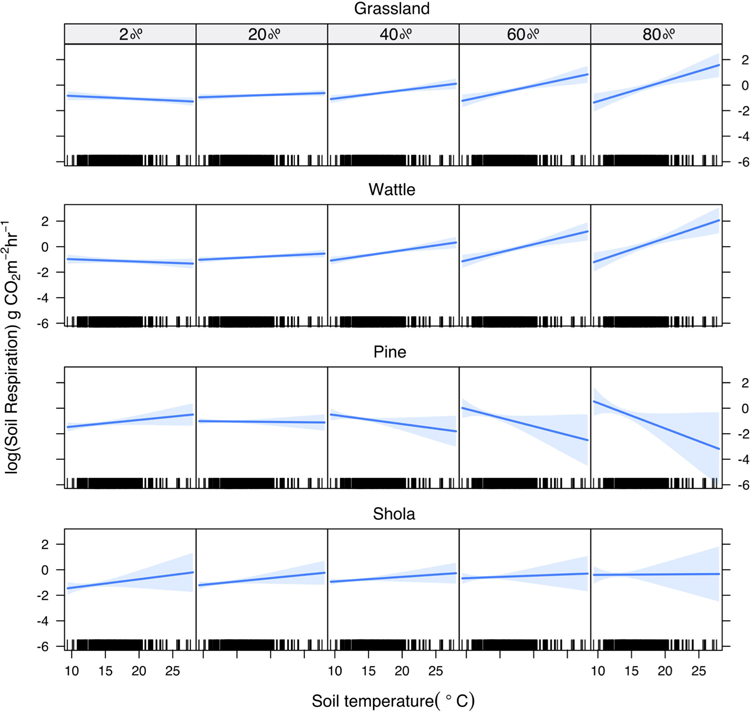
Effect plots depicting the interactive effect of soil moisture and soil temperature across the different land-use types (Each column represent soil moisture in %).

## Discussion

Our results indicate that land-use changes have substantially altered CO_2_ fluxes in this ecosystem. Grasslands consistently had higher CO_2_ effluxes when compared to native shola forests. However, conversion of natural grasslands to alien pine plantations and invasive wattle patches has lowered soil respiration rates. The direction and magnitude of effects on soil respiration due to these land-use changes appear to be collectively governed by the identity of the associated tree species as well as their impacts on local micro-climates. Reductions in soil C effluxes were highest following conversion of grasslands to pine plantations, and somewhat lower following invasion of grasslands by wattle. In addition, conversion of grassland to pine plantations also appears to have altered the sensitivity of soil respiration to changes in soil temperature and moisture.

Soil respiration in all focal land-use categories was primarily governed by precipitation [29–31], peaking during the monsoon when root growth, microbial activity and litter decomposition are typically high [32] and dropping to their lowest levels during the summer. However, intra-annual patterns of precipitation distribution also appear to play an important role in regulating total annual soil CO_2_ effluxes in this system. The highest annual soil respiration levels recorded during 2016-17 in our study (Fig 3) coincided with a more even distribution of precipitation through the year, and a potential lengthening of the growing season, rather than higher overall precipitation during the year.

Across the different land-use types, highest rates of soil respiration rates were recorded in grasslands (Table 1, Fig 3), which is in accordance with previous global syntheses that have reported ~20% higher respiration in grasslands than in comparable forest stands [33]. Respiration rates in shola forests were lower by comparison (Table 1, Fig 3). Invasion of grasslands by wattle reduced soil respiration rates to levels similar to those of natural shola forests while, conversion of grasslands to pine plantations was associated with the greatest reductions in soil respiration (Table 1, Fig 3), consistent with previous large-scale syntheses that report ~10% lower respiration in coniferous stands compared to adjacent broad-leaved forests [32,33]. The low rates of soil respiration recorded in pine plantations is potentially attributable to the poor quality and low nutrient content of pine needles, resulting in slower litter decomposition rates and lower soil respiration rates in pine plantations [32].

Although seasonal patterns of soil respiration were largely governed by soil moisture, instantaneous soil CO_2_ efflux rates were additionally regulated by soil temperature in ways that differed across land-use types. In most land-use types, respiration was largely unaffected, or only marginally increased with soil temperature when moisture levels were low. Low water content is known to limit CO_2_ production in soils, such that temperature effects are typically manifest only when there is sufficient water to allow root and microbial CO_2_ production [29]. However, the nature of temperature-respiration relationships at high moisture levels differed between land-use types (Fig 4). Respiration increased with temperature when moisture was high in both grassland and wattle patches, decreased in the case of pine plantations, and remained largely unchanged in shola forests (Fig 4). The reasons underlying these differences are presently unclear, but are potentially related to differences in the temperature optima of microbial communities in the different land-use types. Given that pine plantations were originally grassland, these results suggest that changes in vegetation type can affect soil C cycling not only by changing above and below-ground C stocks, but also by altering the temperature and moisture sensitivity of respiration even in the absence of any underlying differences in soil type.

## Conclusions

Our results indicate that different vegetation types growing adjacent to one another under the same climatic and environmental conditions can have different soil respiration rates and different sensitivities of soil respiration to temperature and moisture [20]. Soil respiration is a combination of both autotrophic (root) and heterotrophic activity (microbes, soil fauna) [34], and is influenced by several factors including substrate quality and quantity, microbial activity, composition and biomass, and soil temperature and moisture [35 and references therein). All of these likely differ between land-use types, but their relative contributions to the observed differences in soil respiration between land-use types in our study remains unknown. What is clear, however, is that past changes in land use and invasions by exotic species have significantly altered C cycling in this system. Continued invasions and changes in land-use, coupled with changes in temperatures and rainfall climatology are likely to further alter soil respiration in this system in the future. Given the importance of soil respiration for net ecosystem C balance and global carbon budgets, a better understanding of the factors controlling soil respiration in different land-use types is critical in order to predict the long-term responses of C-cycles in this ecosystem.

## Supporting information

## Author contribution

Atul Arvind Joshi: Conceptualization, Methodology, Formal analysis, Investigation, Writing: Original draft

Jayashree Ratnam: Methodology, Writing: review and editing

Harinandanan Paramjyothi: Investigation

Mahesh Sankaran: Conceptualization, Methodology, Formal analysis, Funding acquisition

Supervision, Writing: review and editing

## Acknowledgements

We thank Ministry of Earth Sciences, India (Grant Number: MoES/NERC/16/02/10 PC-11) and National Centre for Biological Sciences, Bangalore for financial support for conducting this study. We are grateful to Tamil Nadu Forest Department for permits. We thank Dr. Jagdish Krishnaswamy (ATREE) and Dr. Ravi Bhalla (FERAL) for sharing precipitation data, and Dr. Kavita Isvaran, Nachiket Kelkar, and Dr. Varun Varma for help with the analysis. We also thank Amol, Kavita, Manaswi, Selvakumar, Susilan, Yadugiri, the FERAL field team in the Nilgiris, residents of Emerald village and the frontline staff of the Nilgiri South Forest Division of Tamil Nadu Forest Department for all their help during field data collection.

## Supporting information caption

**S1 Table.** Linear mixed effect (LME) model of the interactive effect of land-use types, soil moisture and soil temperature on the soil respiration.

