## Supplementary material for "Climate and vegetation collectively drive soil respiration in montane forest-grassland landscapes of the southern Western Ghats, India"

| **Variable** | ***df*** | ***F* - value** | ***p* - value** |
| --- | --- | --- | --- |
| Intercept | 1 | 566.9 | <0.0001 |
| Soil moisture (SM) | 1 | 117.2 | <0.0001 |
| Soil temperature (ST) | 1 | 10.9 | 0.001 |
| Land-use category (L) | 3 | 1.34 | 0.33 |
| SM*ST | 1 | 19.3 | <0.0001 |
| SM*L | 3 | 0.87 | 0.45 |
| ST*L | 3 | 0.11 | 0.95 |
| SM*ST*L | 3 | 4.48 | 0.004 |
